# Temporal and spatial auxin responsive networks in maize primary roots

**DOI:** 10.1101/2022.02.01.478706

**Authors:** Maxwell R. McReynolds, Linkan Dash, Christian Montes, Melissa A. Draves, Michelle G. Lang, Justin W. Walley, Dior R. Kelley

## Abstract

Auxin is a key regulator of root morphogenesis across angiosperms. To better understand auxin regulated networks underlying maize root development we have characterized auxin responsive transcription across two time points (30 and 120 minutes) and four regions of the primary root: the meristematic zone, elongation zone, cortex, and stele. Hundreds of auxin-regulated genes involved in diverse biological processes were quantified in these different root regions. In general, most auxin regulated genes are region unique and are predominantly observed in differentiated tissues compared to the root meristem. Auxin gene regulatory networks (GRNs) were reconstructed with these data to identify key transcription factors that may underlie auxin responses in maize roots. Additionally, Auxin Response Factor (ARF) subnetworks were generated to identify target genes which exhibit tissue or temporal specificity in response to auxin. These networks describe novel molecular connections underlying maize root development and provide a foundation for functional genomic studies in a key crop.

## Introduction

Auxin is a central regulator of root development, playing critical roles in processes such as meristem maintenance and lateral root formation (reviewed in (Atkinson et al., 2014). Root architecture varies among angiosperms and can be influenced by both nutrient and hormone signaling. While maize roots differ considerably in their anatomy and architecture from *Arabidopsis* roots, there are a couple of maize root developmental mutants that have been linked to auxin either directly or indirectly. For example, *rootless undetectable meristem 1* (*rum1*) encodes an AUXIN/INDOLE ACETIC ACID (Aux/IAA) protein that is required for embryonic and postembryonic root formation (von Behrens et al., 2011). In addition, the *rootless concerning crown and seminal roots* (*rtcs*) mutant encodes a LATERAL ORGAN BOUNDARY (LOB) transcription factor which is linked to auxin regulated gene expression (Taramino et al., 2007; Xu et al., 2015). Given that many key auxin signaling components have not been well studied in maize and differential root anatomy may reflect unique regulatory networks, there is a considerable need to increase our knowledge in this area.

The current models for auxin perception and signaling encompass decades of studies (Powers & Strader, 2020). Auxin perception occurs via a co-receptor complex comprised of F-box TRANSPORT INHIBITOR RESPONSE1/AUXIN SIGNALING F-BOX (TIR1/AFB) and Aux/IAA proteins (Calderón Villalobos et al., 2012; Dharmasiri et al., 2005; Kepinski & Leyser, 2004). Because TIR1/AFB proteins encode E3 ubiquitin ligase enzymes, this interaction leads to ubiquitination and degradation of Aux/IAA proteins (Kelley, 2018). Additionally, Aux/IAA proteins (29 family members in *Arabidopsis* and 31 in maize) are transcriptional repressors that actively repress AUXIN RESPONSE FACTOR (ARF) transcription factors in cooperation with TOPLESS co-repressor family of proteins (Szemenyei et al., 2008; Gallavotti et al., 2010). Thus, in the absence of Aux/IAA proteins ARF transcription factors are able to transcriptionally regulate gene expression very rapidly. The ARF protein family (23 members in *Arabidopsis* and 31-36 members in maize) has been recently divided into three classes based on their structure and ability to either activate or repress gene expression (termed “activators” or “repressors”) (Galli et al., 2018). A study by Galli and colleagues uncovered hundreds of auxin responsive genes in maize seedlings that are regulated by particular ARFs. In addition, a recent quantitative genetics study of two maize inbred lines revealed that auxin signaling is a key aspect of maize primary root growth (Wang et al., 2021). Given that auxin responses are known to be influenced by cellular context (Bargmann et al., 2013; Brunoud et al., 2012; Novák et al., 2012; Truskina et al., 2021), an increased resolution of auxin mediated transcription in maize would be beneficial.

Biological networks can describe molecular connections underlying cellular processes and provide insight towards complex phenomena. With respect to auxin signaling, several aspects of auxin action are well-suited to network analyses. For instance, the direct influence of auxin on transcription can be modeled through construction of gene regulatory networks (GRNs). Thus, in this study we performed transcriptome analysis of four regions of the primary root (meristematic zone, elongation zone, cortex, and stele) following 30 and 120 minutes of exogenous indole-3-acetic acid treatment to quantify auxin responsive gene expression with spatial resolution. These data were used to generate novel auxin driven predictive GRNs that underly maize root morphogenesis.

## Methods

### Plant material

Maize seedlings were grown via the rolled towel method as follows. *Zea mays* inbred B73 kernels were surface sterilized in 5% bleach for 15 minutes and rinsed three times with sterile deionized water. For every 10 kernels, three pieces of seed germination paper (Anchor Paper Company, 10×15 L 38# regular weight seed germination paper) were soaked in a solution of freshly prepared Captan fungicide (2.5 g/L). Ten kernels were placed ~ 5 cm from the top of the paper in the middle sheet, covered with the top sheet and rolled into a cylinder lengthwise using the so-called “cigar roll” method. Twelve paper rolls were placed in a 4L Nalgene beaker containing 400 mL of 0.5X Linsmaier and Skoog (LS) pH buffered basal salts (Caisson Labs). The rolled towels were placed in a Percival growth chamber set to 22°C, long day (16 hours light, 8 hours dark) white light at 160 μmol·m^-2^·s^-1^ light intensity. After two days the rolls were opened and the seeds were scored for germination. Any ungerminated kernels were removed and this was designated day 1. After two days of growth the liquid media was poured off and replaced with fresh 400 mL of 0.5X LS. Five days after germination (5 DAG) the seedlings were removed from the towels prior to mock or auxin treatments followed by dissection. Seedlings were placed in 0.5X LS supplemented with 10 *μ*M indole-3-acetic acid (IAA) dissolved in 95% ethanol (“auxin” treatment) or an equivalent volume of 95% ethanol (“mock” control) and incubated at room temperature for 30 and 120 minutes. For each biological replicate, 30-80 primary roots (approximately 2-4 cm in length) were hand dissected into meristematic zone (“MZ”), elongation zone (“EZ”), cortical parenchyma and epidermis (“C”) and stele (“S”) according to previous methods (Saleem et al., 2010; Marcon et al., 2015; Walley et al., 2016) to yield at least 100 mg of tissue per sample and replicate. Total tissue weights per tissue replicate varied from 100 – 600 mg. In total, three biological replicates were collected for each tissue and time point for the transcriptome analysis. Tissues were immediately flash frozen in liquid nitrogen and stored at −80 until all replicates were harvested.

### RNA extraction and transcriptome sequencing

Root tissues were ground to fine powder in liquid nitrogen using a pre-chilled mortar and pestle. RNA was extracted using a modified Trizol/RNeasy hybrid protocol (Walley et al., 2010) with Trizol reagent (Invitrogen) and a Zymo Direct-zol RNA miniprep kit (Zymo). RNA concentrations and quality were initially checked using a Nanodrop and Qubit. RNA was then submitted to the Iowa State University (ISU) DNA facility. Submitted RNA samples were quality checked at the ISU DNA facility via Bioanalyzer and then used to generate QuantSeq 3’mRNA libraries using a Lexogen 3’ mRNA-Seq FWD kit and 48 unique indices (Moll et al., 2014). Libraries were run on an Illumina HiSeq 3000 to generate 100 bp single-end (SE) reads. Raw data files obtained from the ISU DNA facility were stored on the Large Scale Storage (LSS) at ISU.

### RNA-seq analysis

Raw sequence files were deposited at the NCBI Sequence Read Archive (BioProject accession number PRJNA791716). Files were checked via FASTQC to obtain sequence quality information. The fastq files were then ran through the following bioinformatic mapping pipeline as suggested by Lexogen. First, adapters and low-quality tails were removed in *bbduk* from the BBTools suite (sourceforge.net/projects/bbmap/). Alignment to the B73 v4 genome (Jiao et al., 2017) was performed using STAR (Dobin et al., 2013). Indexing was performed using SAMTools (H. Li et al., 2009). HTSeq (Anders et al., 2015) was used to generate count files which were then analyzed via PoissonSeq (J. Li et al., 2012). Differentially expressed genes were identified using a false discovery rate (FDR; adjusted p-value) of q-value <0.1.

### Gene regulatory network analysis

Transcription factor (TF)-centered gene regulatory networks (GRNs) were generated using SC-ION version 2.1 (Clark et al., 2021) and annotated maize TFs from Grassius (Yilmaz et al., 2009). We first clustered the transcript data by root region (MZ, EZ, C, or S) using Independent Component Analysis (ICA) (Nascimento et al., 2017) implemented in SC-ION. We then input the TMM-normalized counts matrix from our RNA-seq analysis, coupled with “regulator” (DE TFs only) and “target” (all DE genes) lists. Those input files were then used to analyze each ICA cluster by the SC-ION adapted version of the GENIE3 (Huynh-Thu et al., 2010) network inference algorithm, which output a table with the predicted regulator-target interactions as well as a numeric “weight” value for each pair indicating the confidence of their connection. The SC-ION generated GRNs were imported into Cytoscape (Shannon et al., 2003) for visualization.

### Other software

UpSet plots were generated using the UpsetR package in RStudio and ordered by frequency. Additional code was written to extract gene identifiers among shared lists of differentially expressed genes within an UpSet plot; data processing scripts are available from a github repository: https://github.com/mmcreyno92/AuxinRootAtlas. Gene ontology (GO) enrichment analysis was performed in PANTHER using the Zea mays B73 reference genome with a Fisher’s Exact test type and a false discovery rate correction. Enriched GO terms were plotted in R using GO R code (Bonnot et al., 2019).

## Results and Discussion

### Auxin responsive transcriptome profiles in primary maize roots across space and time

The overall goal of this study was to characterize the auxin responsive transcriptome within maize primary roots across four key cellular regions: the meristematic zone (MZ), elongation zone (EZ), and the cortex (C) and stele (S) within the differentiation zone (Marcon et al., 2015; Paschold et al., 2014; Walley et al., 2016). Five-day-old B73 maize primary roots were treated with 10 μM indole-3-acetic acid (IAA), hereafter referred to as “auxin” or an equivalent volume of 95% ethanol “mock” control for 30 and 120 min and then dissected into four regions (Figure 1A). Transcriptome profiling was performed on these tissues using the 3’ QuantSeq method (Moll et al., 2014) with three biological replicates for each tissue/treatment/time. From this analysis we identified 32,832 transcripts in total across all tissues. Within the meristematic zone, relatively few auxin responsive genes were observed (Figure 1B). In contrast, hundreds of genes were induced or repressed following auxin treatment within the elongation zone, cortex, and stele. This result suggests that meristematic zone cells may be less sensitive to exogenous auxin effects and/or have altered cellular states which buffer transcriptional responses. For example, meristematic zone cells could contain a larger proportion of heterochromatin compared to neighboring differentiated cells within the root.

**Fig. 1.**
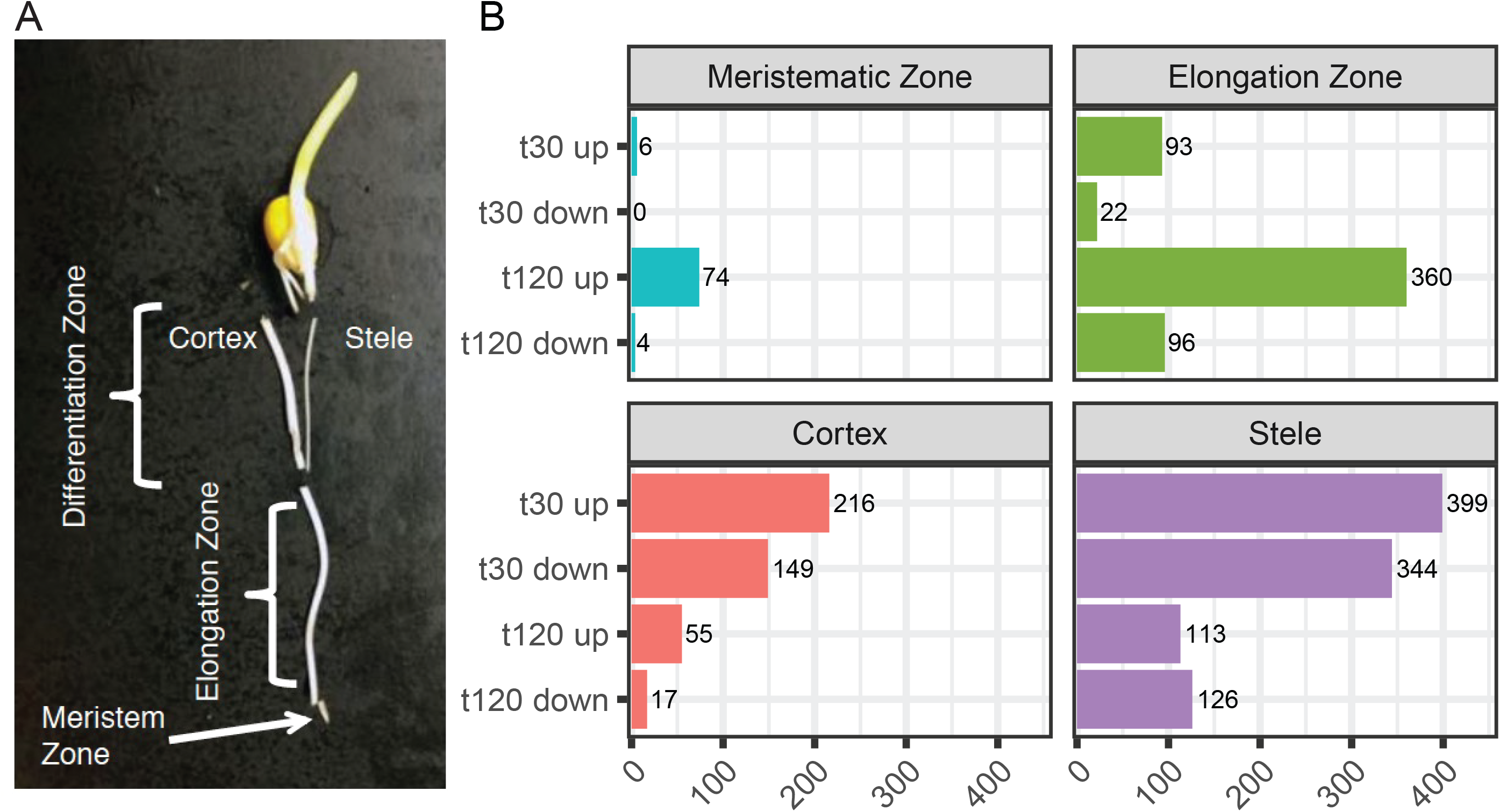
Identification of auxin responsive genes across four key regions of the primary maize root. (A) Light micrograph of maize primary dissected root regions profiled in this study. Five-day-old primary maize roots were dissected into the four regions indicated. The distal 2 mm of the root tip corresponds to the meristematic zone (MZ). The elongation zone (EZ) is the proximal zone adjacent to the MZ root tip up to where the root hairs emerge. The differentiation zone, starting with the root hair zone, was mechanically separated into cortex (C) and stele (S) by snapping the root from the kernel and pulling the stele out from the cortex. (B) Differentially expressed genes within each root region at 30 min (t30) and 120 min (t120) were identified by comparing auxin treated samples to mock treated samples at q<0.1.

Within the cortex and stele more auxin regulated genes are observed at 30 min compared to 120 min. In contrast, within the elongation zone, we observed more up-regulated genes at 120 min compared to 30 min. These patterns may reflect the altered chromatin state or transcription factor properties associated with cellular state as cortex and stele cells are further differentiated compared to elongation zone cells.

### Auxin regulated gene expression in maize primary roots is region specific

Auxin mediated gene expression is context dependent. To examine shared and uniquely regulated transcripts across the four sampled root regions we generated UpSet plots comparing auxin responsive transcripts within each time point (Figure 2) as well as differentially expressed (DE) between tissues (Supplemental Figure 1). Given that there are numerous possible comparisons with four regions and two categories of DE (up or down) we selected the top 20 comparisons for visualization. At both 30 and 120 minutes after auxin treatment, relative to mock treatment, the majority of the observed DE genes are region specific. At 30 minutes after treatment auxin up- and down-regulated genes across regions include both concordant and discordant properties. In contrast, at 120 minutes after treatment the observed DE genes in common between tissues are only concordant. For example, at 30 min there are 22 transcripts which are up-regulated in the cortex but repressed (down) in the stele. This result suggests that early auxin-mediated transcriptional changes may include both repression and activation at the same genes in a tissue specific manner, while later effects (i.e. 2 hours) of auxin may uniformly influence suites of genes irrespective of cellular context.

**Fig. 2.**
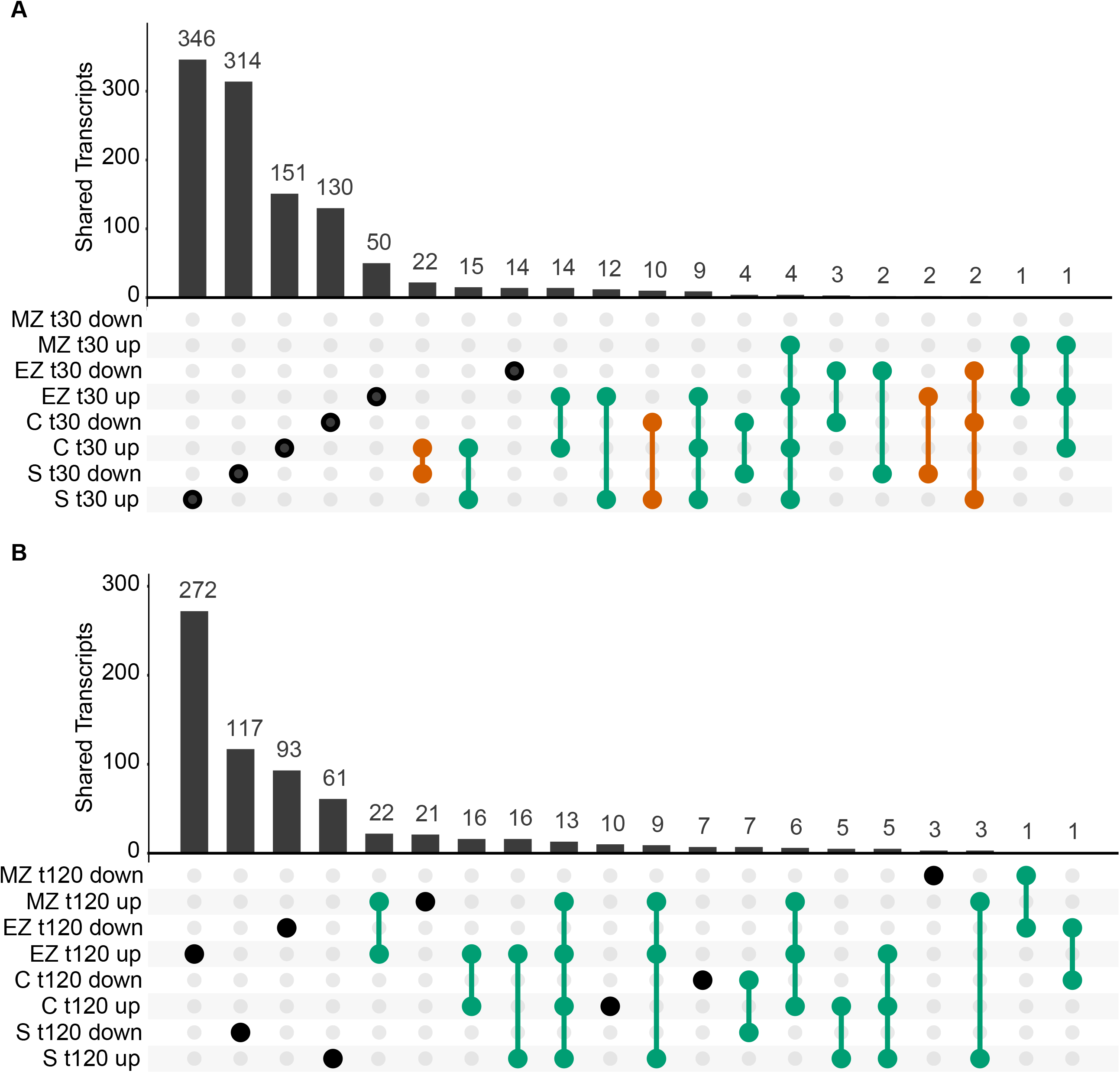
A comparison of differentially expressed genes in maize roots across four regions at two different timepoints in response to auxin. (A) UpSet plot of differentially expressed transcripts at 30 min (t30). (B) UpSet plot of auxin-responsive DE transcripts at 120 min (t120). Concordant and discordant comparisons are indicated in green and vermillion, respectively. Abbreviations used: MZ = meristematic zone, EZ = elongation zone, C = cortex, S = stele, auxin = indole-3-acetic acid treatment compared to mock treatment.

These comparisons provide the opportunity to identify robust auxin responsive transcripts which are up-regulated irrespective of tissue or time point. For example, there are four transcripts which are auxin induced in the meristematic zone, elongation zone, cortex, and stele at 30 min and 13 such transcripts at 120 min. These transcripts include *AUX/IAA-transcription factor 22* (*IAA22/Zm00001d013707*), *Aux/IAA24* (*Zm00001d018414*), *DIOXYGENASE FOR AUXIN OXIDATION 1* (*DAO1/Zm00001d003311*) and *AUXIN AMIDO SYNTHETASE2* (*AAS2/Zm00001d006753*). Notably, these are all encoded by genes with annotated functions in auxin response and auxin metabolism and thus may reflect pathway feedback. From this analysis a set of auxin responsive marker genes have now been identified which can facilitate future studies on auxin signaling in maize roots.

### Distinct biological processes are enriched among auxin regulated genes across root regions

To determine if particular biological processes are auxin-regulated in root regions we performed a gene ontology (GO) enrichment analysis. Many of the observed enriched GO terms are congruent with a previous study that examined the transcriptome profile of these root zones in the absence of treatment (Paschold et al., 2014), but we also identified a number of novel GO terms associated with hormone signaling, cell cycle, and gene regulation (Figure 3, Supplemental Figure 2, Supplemental Table 2). In general, auxin induced genes are associated with transcription, auxin-activated signaling pathway, and gibberellin metabolism. In contrast, auxin repressed genes are associated with cell cycle, cell division, and chromatin silencing.

**Fig. 3.**
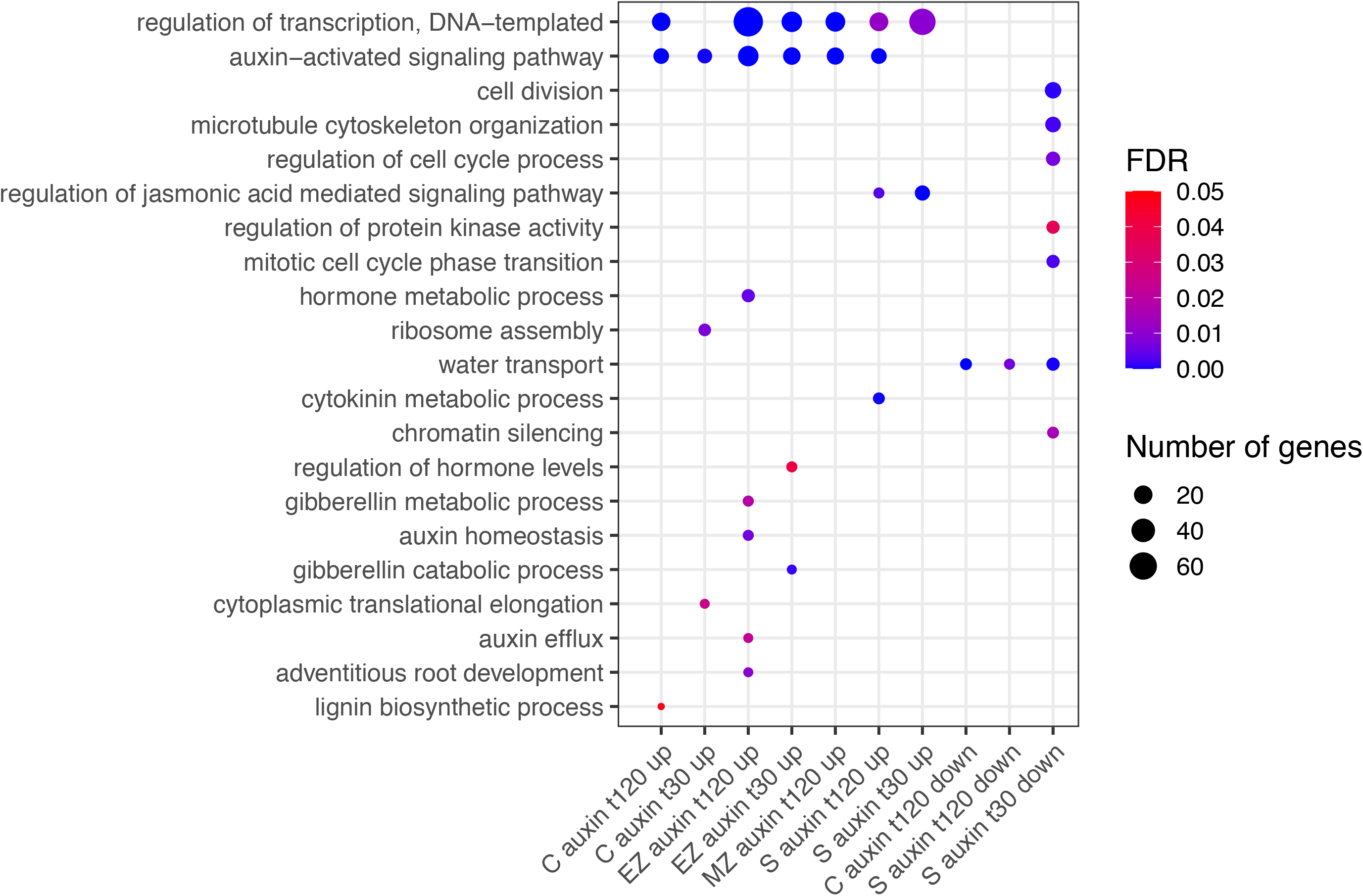
Auxin responsive genes between root regions are enriched in several gene ontology (GO) terms related to biological processes. Significant GO terms of interest in auxin down-regulated genes (“down”) and auxin up-regulated genes (“up”) are indicated on the y-axis. False discovery rate (FDR) is color-coded from blue (0.00) to red (0.05). Size of the dot indicates the number of enriched genes within each GO term. Abbreviations used: MZ = meristematic zone, EZ = elongation zone, C = cortex, S = stele, t30 = 30 min, t120 = 120 min.

In addition, we examined GO term enrichment between root regions to uncover tissue-specific processes that may underlie root structure (Supplemental Figure 2). In general, most GO terms appear to be tissue specific and many of the observed enriched GO terms are congruent with the previous study (Paschold et al., 2014). A couple enriched GO terms standout among the many observed. First, transcripts involved in protein phosphorylation are more abundant in the stele compared to the meristem or the neighboring cortex. Another GO term observed across several tissue comparisons is ‘microtubule-based movement’, which is to be expected for cells undergoing cell elongation and/or differentiation. Secondary cell wall biogenesis is more prevalent in elongation zone expressed genes compared to meristem zone transcripts, which fits with our current understanding of cell wall composition across the primary root. Altogether these results support the notion that root tissues exhibit unique cellular processes that may be linked to function.

### Identification of spatially distinct auxin gene regulatory networks within primary roots

In order to infer regulatory relationships between auxin responsive root transcription factors (TFs) and their targets we generated a gene regulatory network (GRN). To reconstruct the predictive GRN we implemented our network inference pipeline, SC-ION, which is an extension of RTP-STAR and has been shown previously to successfully identify novel TF roles in response to hormone treatment (Clark et al., 2019; Broeck et al., 2021; Clark et al., 2021). The resulting GRN consisted of 15,856 nodes (genes) with a total of 86,461 directed edges (Figure 4 and Supplemental Table 3). A circular layout visualization of the complete GRN illustrates the presence of several distinct groups (circles) based on their underlying tissue enrichment, either within a singular root region (MZ, EZ, C, or S) or between multiple combinatorial root zones (e.g. MZ+EZ, MZ+EZ+C, etc.) based on the SC-ION generated independent component analysis (ICA) (Nascimento et al., 2017) clustering assignments (Supplemental Table 3).

**Fig. 4.**
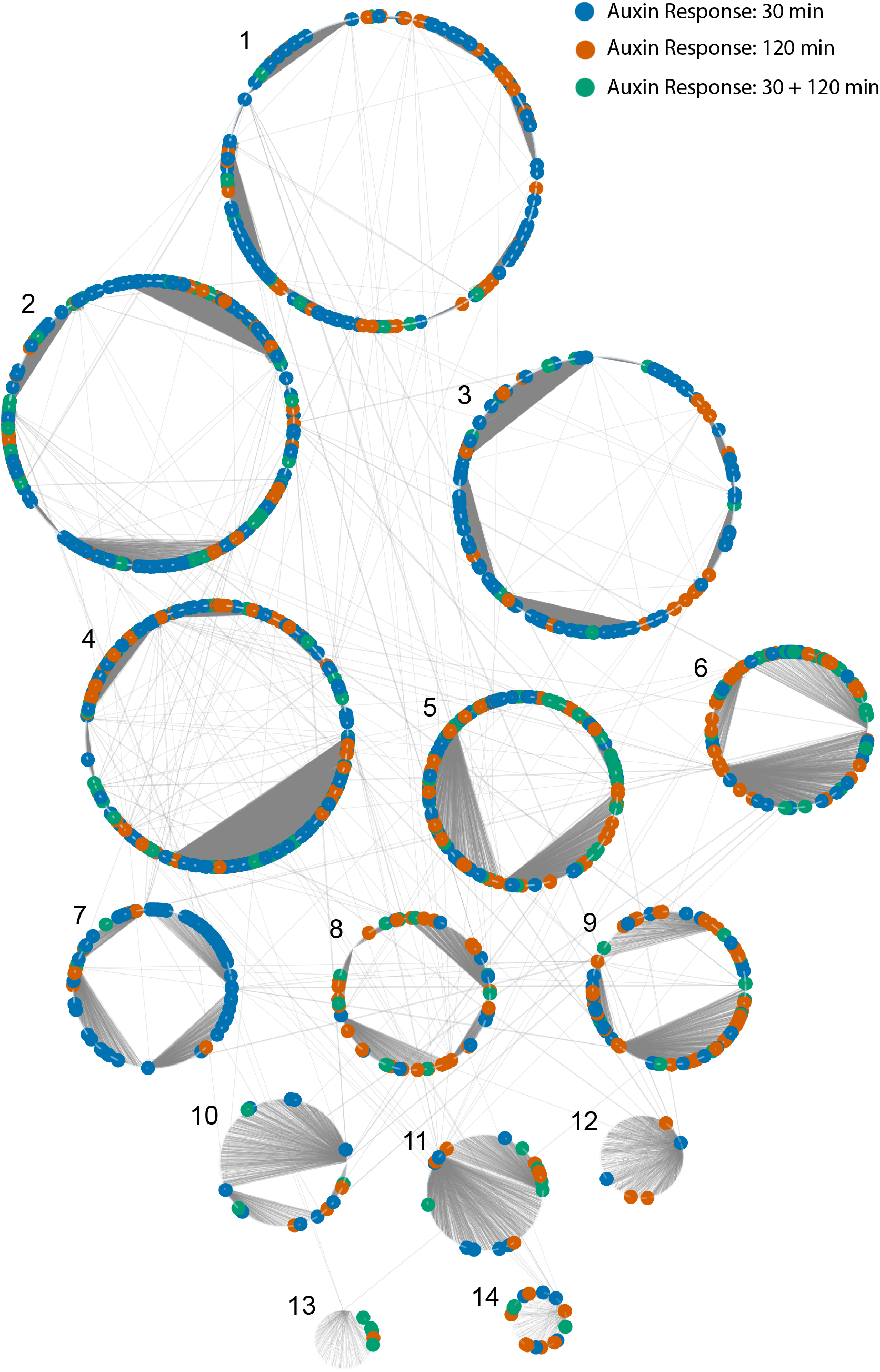
A spatiotemportal auxin responsive gene regulatory network in maize primary roots. The nodes (genes) are arranged in numbered circles represent groupings of nodes (genes) that clustered together and were enriched within the same tissues. Colored nodes represent genes that are differentially expressed following auxin treatment. The temporal response to auxin is indicated by node color: blue = auxin responsive at 30 min, vermillion = auxin responsive at 120 min, and green = auxin responsive at both time points sampled. Each circular node represents a distinct cluster based on tissue: 1 = C + S, 2 = C, 3 = MZ, 4 = S, 5 = EZ, 6 = EZ + S, 7 = MZ + S, 8 = MZ + EZ, 9 = EZ + C, 10 = MZ + C, 11 = EZ + C + S, 12 = MZ + C + S, 13 = MZ + EZ + S, 14 = MZ + EZ + C.

The nodes contained within our GRN fell into 1 of 14 tissue enrichment groups with content sizes ranging from 2,862 nodes enriched in the C+S (group 1) down to 98 nodes in the MZ+EZ+C (group 14). The tissue enrichment groupings also featured varying numbers of auxin responsive genes and a high degree interconnectedness as evidenced by the number of edges linking nodes within a grouping. Using predicted regulator data from the GRN, we investigated the regulatory relationships of genes known to be involved in auxin signaling and maize root architecture (Figure 4 and Supplemental Table 3). One such transcription factor of interest, RTCS1 (Zm00001d027679), was predicted in the GRN to regulate 57 target genes including the auxin responsive gene AUX/IAA32 (Zm00001d018973). Additional root development associated transcription factors represented in the data include other LBD-transcription factor family members along with multiple members of the maize SHI/STY (SRS) family (Gomariz-Fernández et al., 2017), including a known transcriptional activator *lateral root primordia 1* (Zm00001d011843) that is required for maize root morphogenesis (Zhang et al., 2015).

ARF transcription factors represent a critical regulatory component of the auxin response, thus we set out to inspect their target gene relationships within the GRN at a deeper level. First, we identified all of the annotated ARFs present in the GRN and observed that 27 of the 33 expressed maize ARF family members were present in the GRN. For these 27 ARFs and their first node neighbor targets, we generated subnetworks that were visualized in prefuse force directed layout in Cytoscape (Figure 5 and Supplemental Table 4). Notably, representative ARFs from each of the four distinct evolutionary ARF clades (Galli et al., 2018) were found to be present in the GRN, including 12 clade A ARFs, 6 clade B ARFs, 4 clade C ARFs, and 5 ETTIN-like ARFs (visualized as pink nodes in Figure 5). In general, most ARFs had target nodes that were DE in response to auxin (coded blue and orange in Figure 5) and exhibit unique targets compared to one another. In three instances there are several ARFs that have shared target genes with one another, including ARF18 and ARF7; ARF8, ARF23 and ARF25; and ARF24 and ARF36. Notably, the ARFs with shared targets span phylogenetic clades, suggesting that properties of ARFs cannot be predicted based on sequence evolution alone. In addition, we examined the ARF target genes and found that they include auxin-related genes belonging to the *ARF* (5), *Aux/IAA* (8), *SAUR* (3), and *GRETCHEN HAGEN* (*GH3*) (1) families. Such targets are well-known types of auxin responsive genes (Bargmann et al., 2013; Galli et al., 2018; Lewis et al., 2013; Nemhauser et al., 2006) and indicate that ARF proteins are engaged in feedback loops.

**Fig. 5.**
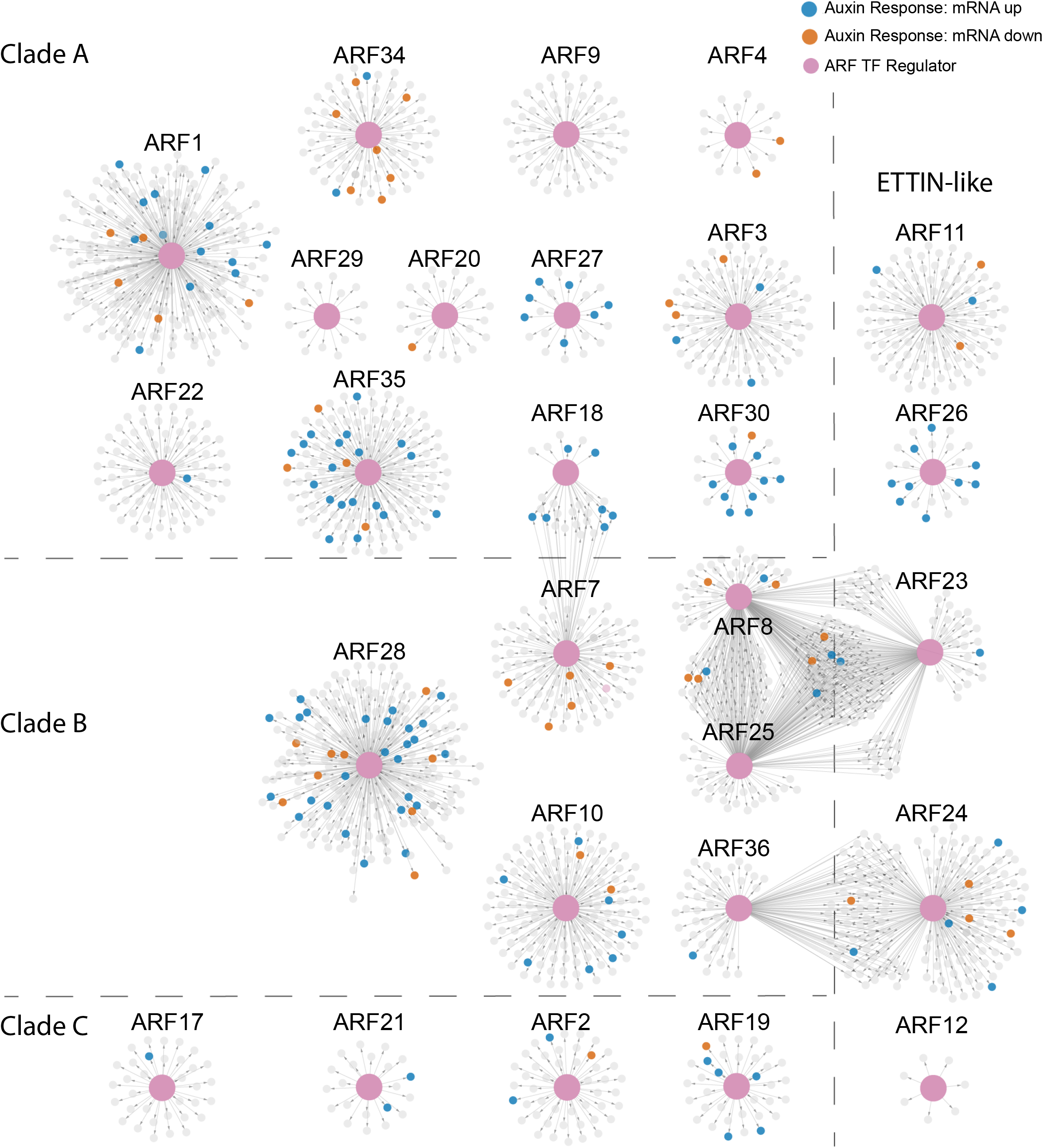
Auxin Response Factor (ARF) transcription factor gene regulatory subnetworks associated with primary maize roots. The networks are arranged by clade classification: Clade A, Clade B, Clade C, or ETTIN-Like. The central enlarged pink nodes within each network represent the ARF of interest labelled above the network and the connected small nodes represent that ARFs target genes. Target genes are colored according to the directionality of their transcript expression in response to auxin: grey = no significant transcript change, vermillion = decreased transcript level, blue = increased transcript level.

In this work we utilized a combinatorial approach of transcriptome analysis and gene network inference to identify temporally auxin responsive genes across root tissue types. By elucidating the complex inner workings of auxin-mediated gene expression during primary maize root development we can begin to answer questions surrounding root architecture in this key crop. A molecular understanding the dynamics of root growth can aid in informing strategies to create next-generation crops with more efficient water and nutrient uptake capabilities.

## Acknowledgements

We wish to thank Diana Burkart for her assistance with Lexogen QuantSeq data analysis and Natalie Clark for her assistance with SCION. This work was supported by USDA NIFA AFRI Predoctoral Fellowship to MRM (Award No. 2021-67034-35188), USDA NIFA AFRI grant to DRK and JWW (Award No. 2020-67013-30914), start-up funds to DRK from Iowa State University (ISU), funding from the ISU Plant Science Institute to JWW, and Hatch Act and State of Iowa funds to DRK (Project No. IOW03649) and JWW (Project No. IOW04108).

## Supplementary material

**Supplemental Figure 1.** UpSet plot comparing differentially expressed genes across maize root regions in “mock” treated samples. Abbreviations used: C = cortex, S = stele, EZ = elongation zone, MZ = meristem zone, up = up-regulated genes, down = down-regulated genes. Concordant gene expression differences are indicated in bluish green while discordant gene expression differences are in vermillion.

**Supplemental Figure 2.** Gene ontology (GO) terms enriched in differentially expressed genes between root regions profiled. Comparisons are indicated such that control condition is first and the control is second. For example, “S/C up” means that transcripts up in the stele (S) relative to the cortex (C) are enriched for the indicated GO terms. Abbreviations used: FDR = false discovery rate, C = cortex, S = stele, EZ = elongation zone, MZ = meristem zone, up = up-regulated genes in both tissues, down = down-regulated genes in both tissues. FDR values are colored according to the heatmap shown, going from blue to red.

**Supplemental Table 1.** Differential expression analysis data from QuantSeq 3’ mRNA-Seq. Workbook contains TMM-normalized expression values for all samples analysed with each additional sheet containing statistical analysis values for each pairwise comparison generated by PoissonSeq.

**Supplemental Table 2.** Gene Ontology (GO) enrichment analysis data of comparisons within tissue mock +IAA treated samples as well as between tissue samples.

**Supplemental Table 3.** Full gene regulatory network construction data. Independent Component Analysis (ICA) clustering assignments, SC-ION adapted GENIE3 output table, assigned tissue enrichment by cluster, and Cytoscape node table export data.

**Supplemental Table 4.** Auxin Response Factor (ARF) subnetwork data. SC-ION adapted GENIE3 output table and Cytoscape node table export data.

